# Epigenomic and transcriptomic differences between low-grade and acute inflammation in LPS-induced murine immune cells

**DOI:** 10.1101/2021.03.28.437377

**Authors:** Lynette B. Naler, Yuan-Pang Hsieh, Shuo Geng, Zirui Zhou, Liwu Li, Chang Lu

**Affiliations:** Department of Chemical Engineering, Virginia Tech, Blacksburg, Virginia, USA; Department of Biological Sciences, Virginia Tech, Blacksburg, Virginia, USA

## Abstract

Chronic, low-grade inflammation has a widespread and significant impact on health, especially in Western society. While inflammation is beneficial for the removal of microbes, low-grade inflammation never resolves and can cause or worsen other diseases. The process by which low-grade inflammation occurs remains poorly understood. Here we exposed murine bone-marrow derived monocytes to chronic lipopolysaccharide (LPS) stimulation at low dose or high dose, as well as a PBS control. The cells were profiled for genome-wide H3K27ac modification and gene expression. The gene expression of TRAM-deficient and IRAK-M-deficient monocytes with LPS exposure was also analyzed. We discover that low-grade inflammation preferentially utilizes the TRAM/TRIF-dependent pathway of TLR4 signaling, and induces the expression of interferon response genes. In contrast, acute inflammation uniquely upregulates metabolic and proliferative pathways that also appear to be TRAM-dependent. The extensive differences in the epigenomic landscape between low-dose and high-dose conditions suggest the importance of epigenetic regulations in driving differential responses. Our data provide potential targets for future mechanistic or therapeutic studies.

## Background

Low-grade, or chronic inflammation, plays a large role in the onset or exacerbation of many mental and physical diseases, including Alzheimer’s disease^1^, depression^2^, diabetes^3^, and some cancers^4^. Despite this, there still are no canonical markers for low-grade inflammation, partly due to a correlation of inflammatory activity with age^5,6^. Some recent studies in humans have attempted more sophisticated integrated approaches that show promise, but more research is required to identify key biomarkers and how they should be interpreted^6–8^.

Low-grade inflammation is often associated with certain lifestyle choices, like Western-style eating practices^9^, sedentary behavior^10,11^, sleep deprivation^12^, and social stress^13^. In fact, individuals who underwent major childhood stressors have elevated mortality and morbidity of immune or chronic diseases later in life, and epigenetic programming has been proposed^14^. This is further supported by twin studies that showed most variation in the immune system is due to non-heritable (e.g. epigenetic) influences^15^.

In a bacterial infection, components of the pathogen^16^ are recognized by pattern-recognition receptors (PRRs) that are found on immune and non-immune cells^17–20^. This, in turn, leads to a signaling cascade that is used to recruit immune cells, such as neutrophils and monocytes/macrophages, which ingest the microbes or release antimicrobials^21,22^. Once the threat is eliminated, pro-inflammatory macrophages uptake spent neutrophils, which reprogram the macrophages to an anti-inflammatory state^23,24^. Anti-inflammatory macrophages help promote healing by protecting against tissue damage, clearing out debris, and producing growth factors^24^. However, when macrophage and monocyte inflammatory activity is not resolved then it can lead to low-grade inflammation and chronic diseases^25–30^.

Lipopolysaccharide (LPS) is an endotoxin and component of the cell wall of Gram-negative bacteria, such as *Escherichia coli*, that also has been found at low levels in individuals who have chronic diseases or detrimental lifestyle choices as mentioned previously^31,32^. It has been used to study inflammation both *in vivo* and *in vitro*, although more often with high-doses of LPS. While persistent low-dose LPS has been shown to lead to a low-grade inflammatory state, the mechanism by which it does so is still not entirely known, though could be due to sustained activity of the inflammatory NF-κB^32–35^.

LPS stimulation activates two different pathways, which both have pro- and anti-inflammatory aspects. First, LPS binds to Toll-like receptor 4 (TLR4) and activates the MyD88-dependent pathway^32,36,37^. The MyD88-dependent signaling cascade activates mitogen-activated protein kinases (MAPKs), NF-κB translocation to the nucleus, and transcription factors CREB and AP-1, as well as induces expression of pro-inflammatory genes such as TNF-α, IL-6, and COX-2^37^. However, it is also involved in the production of anti-inflammatory cytokines like IL-10. In addition, MyD88-dependent signaling leads to an increase in glycolysis, synthesis of acetyl-CoA, and synthesis of fatty acids. Next, TLR4 is endocytosed, ending the MyD88-dependent pathway and triggering the TRIF-dependent pathway^37^. The TRIF-dependent pathway activates IRF3 and IRF7 to induce expression of type I interferons (IFN), CCL5, and CXCL10. It is also involved in antiinflammatory cytokine IL-10 production and is important for TNF-α expression^37,38^. Furthermore, in macrophages, the TRIF-dependent pathway also upregulates cellsurface CD40, CD80, and CD86 which are necessary for antigen presentation for T lymphocytes, bridging the gap between the innate and adaptive immune system^37,39^. Finally, both pathways are involved in the canonical activation of the NRLP3 inflammasome, which is responsible for the activation of the inflammatory cytokine, IL-1β, and can cause pyroptosis, a form of programmed cell-death^37^.

The epigenome plays a large role in the behavior and identity of macrophages. For example, the epigenome of tissue-resident macrophages is affected by their local microenvironment and they can even be reprogrammed by transplanting them to a different location^40^. They also have varied transcriptional signatures during efferocytosis, a process where apoptotic cells are cleared^41^. LPS-stimulation has also been shown to have a significant, and sometimes lasting, effect on the histone modifications of macrophages^42^. LPS-induced tolerance affects the epigenome of macrophages by inhibiting induction of inflammatory genes while leaving antimicrobial genes unaffected^43^. Research into means of targeting and altering the epigenome of inflammatory macrophages into an anti-inflammatory state has also increased in recent years, with studies showing possible therapeutic potential^44^. Despite this, there is little research on how low-doses of LPS affect the epigenome^45–48^ or transcriptome^48–52^ of monocytes or macrophages differently compared to high-doses of LPS.

In this study, we profiled the histone mark H3K27ac and performed RNA-seq analysis of murine bone marrow-derived monocytes that are exposed to low and high-dose levels of LPS, as would be seen in low-grade and acute inflammation, respectively. Low-input technologies including Microfluidic Oscillatory Washing ChIP-seq (MOWChIP-seq^53,54^) and Smart-seq2^55,56^ were used for the epigenomic and transcriptomic profiling, respectively. We compared the conditions to extract the effects of LPS-dosage on the epigenome and, in turn, the transcriptome of immune cells. Furthermore, we also analyzed the effects of TRAM deletion, necessary for the TRIF-dependent pathway^37^, and IRAK-M deletion, which suppresses TLR-signaling and is associated with endotoxin tolerance^57^. Altogether, this analysis could help elucidate the means by which subclinical concentrations of endotoxin can lead to low-grade inflammation as opposed to acute inflammation.

## Results

Bone marrow-derived monocytes (BMDMs) were isolated from mice and dosed with PBS, low-dose LPS (100 pg/mL), or high-dose LPS (1 μg/mL). We profiled H3K27ac using MOWChIP-seq^58^ and performed RNA-seq with two replicates per condition (Suppl. Tables S1 and S2). We found very high average correlation between ChIP-seq replicates of 0.986 (Fig. 1a). Between conditions, the highest correlation is between PBS and low-dose (0.957) and the lowest correlation between PBS and high-dose (0.896), with low-dose and high-dose falling in-between (0.939). This suggests that the increasing LPS dosage causes a concomitant change to H3K27ac signal. When looking at genome-wide H3K27ac signal, we see that increasing the dose of LPS tends to reduce the H3K27ac signal (Fig. 1b). In fact, the overall H3K27ac signal at peaks also tends to go down with increasing LPS-dosage (Fig. 1c). Furthermore, the number of peaks present in each sample as decreased with increasing dosage (PBS = 25,679, Low = 21,597, High = 14,659). As such, many of the peaks that were present in PBS samples were not present in Low-dose samples (6,566) and even more (12,602) were not present in High-dose (Fig. 1d). However, Low-dose samples gained a small number of peaks (2,484) while High-dose samples gained even fewer (1,582). In addition, the fraction of peaks near promoters increases with increasing LPS dosage, yet, this primarily was due to a reduction of distal peaks, rather than an increase of proximal peaks (Fig. 1e). However, some of the peaks gained by Low-dose and High-dose conditions were proximal to gene promoters. Since H3K27ac also localizes to active promoters, this suggests that LPS dosage is activating genes that are not active under normal conditions^59^.

**Figure 1.**
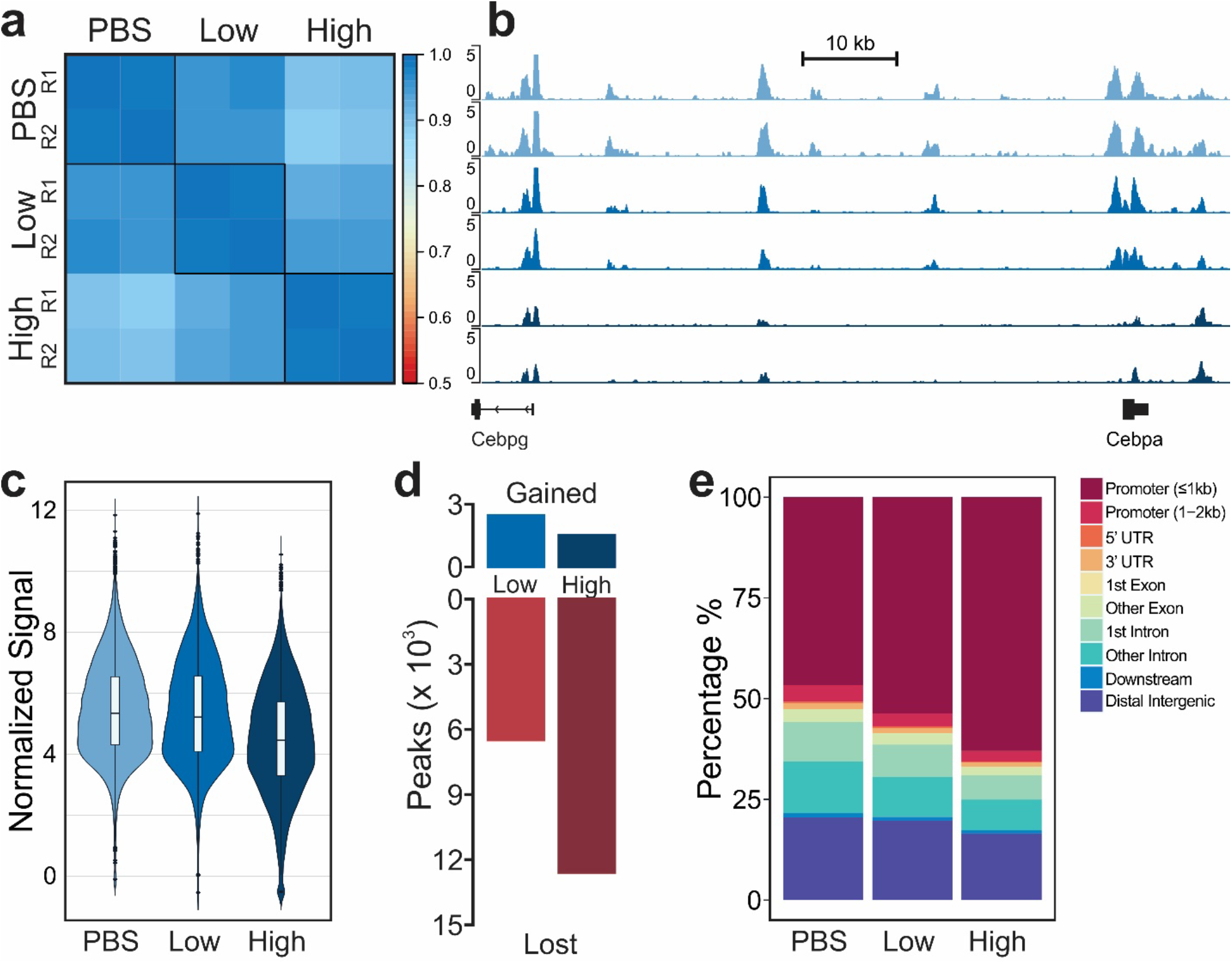
Overview of epigenomic data for murine BMDMs dosed with PBS, low-dose LPS, or high-dose LPS. (a) Pearson’s correlation of normalized H3K27ac signal around promoter regions (TSS +/− 2 kb). (b) Tracks of normalized H3K27ac signal for replicates dosed with PBS, low-dose LPS, or high-dose LPS (from top to bottom). (c) Distribution of normalized H3K27ac signal at peaks. (d) Number of peaks gained or lost in low-dose or high-dose conditions compared to PBS. (e) Percentage of peaks at genomic locations.

Although H3K27ac does mark active transcription, it plays a more pivotal role in long-range gene regulation at enhancer regions. Therefore, we determined the locations of enhancers, which were defined as H3K27ac^high^ regions that do not overlap with regions near transcription start sites (TSS) and can be linked to genes in conjunction with RNA-seq data (see Methods). The number of enhancers also decreased with increasing LPS-dosage (PBS=5,738, Low=4,400, High=2,711). Normalized H3K27ac signal for the conditions at each enhancer was clustered with k-means clustering (Fig. 2a). The genes linked to the enhancers in each of the clusters were analyzed for overrepresentation of Gene Ontology biological process gene sets (Fig. 2b, Suppl. Tables S3 and S4). We then separated these clusters into three groups: dosage correlated (I, II, III), acute inflammation (IV and V) and low-grade inflammation (VIII).

**Figure 2.**
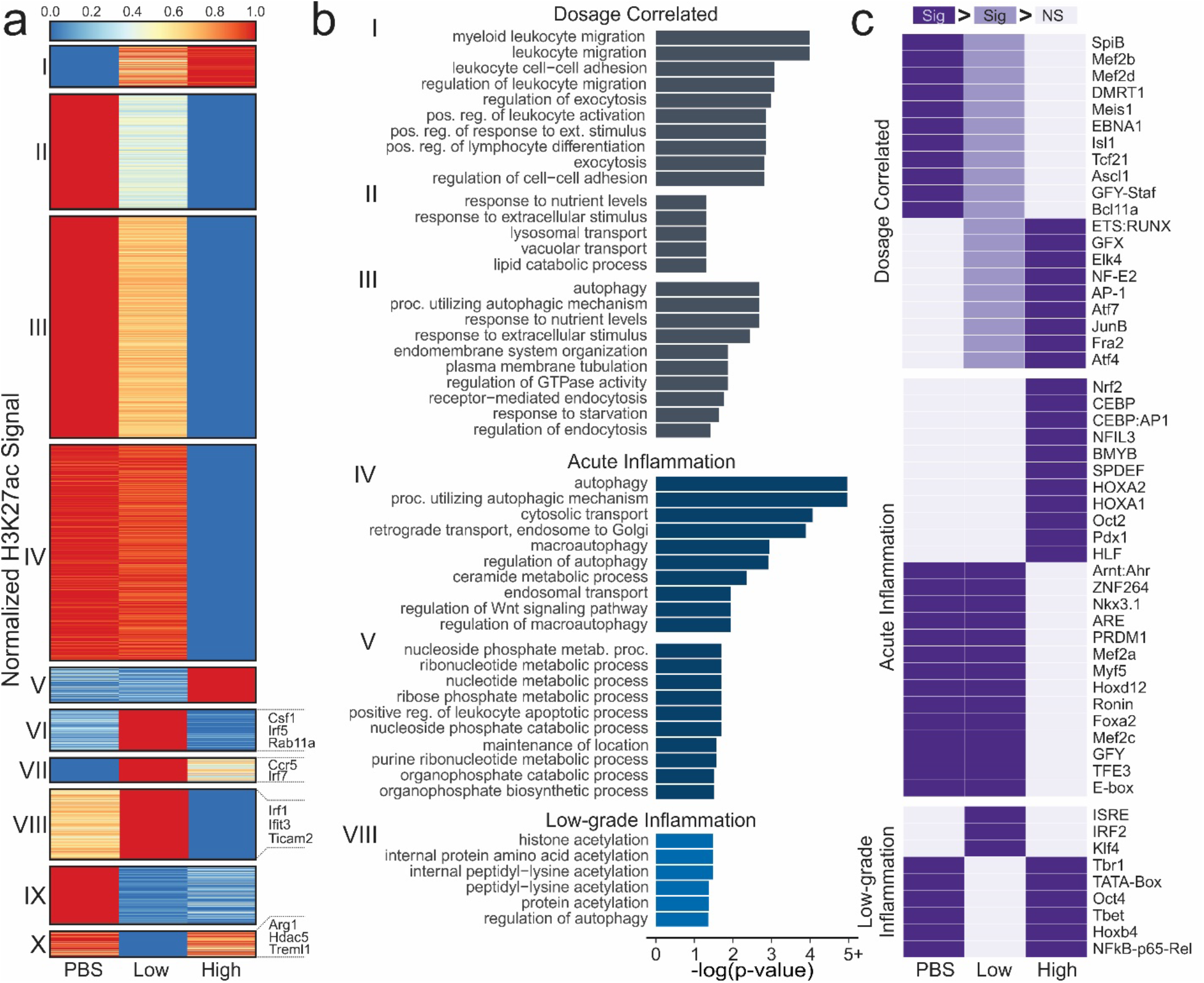
Effect of LPS dosage on enhancers. (a) K-means clustering of enhancers present in PBS, Low-dose, or High-dose samples (color scale is weird) (b) Significant (FDR < 0.05) Gene Ontology biological process gene sets for clusters from (a). (c) Significantly enriched motifs (p < 1 × 10^−6^) that are different in two comparisons. One purple cell denotes enrichment over the two gray cells and two denotes enrichment over the third cell. One light and one dark purple signifies dark is enriched over light, and light is enriched over gray.

The dosage correlated group, including clusters I, II, and III, had either increased or decreased H3K27ac signal with increased LPS-dosage. Most obviously, we see that Cluster I, which increases in expression from PBS to High-dose, is filled with leukocyte-related (which include monocytes and macrophages) gene ontologies, which play a key role in inflammation^60,61^. As such, it is reasonable that with increased LPS-dosage we would see a stronger inflammatory response. In Clusters II and III, which decreased from PBS to High-dose, we see several pathways related to autophagy and endocytosis. It has been shown in literature that LPS-stimulation induces autophagy as a means of mediating the inflammatory response^62–65^. Upon further investigation into the enhancer-linked genes in these autophagy-related processes, we find genes such as Trem2^66^, Sesn1^67^, and Nrbf2^68^, all of which inhibit the inflammatory effect of macrophages. Differential H3K27ac expression at gene promoters also shows that increased LPS-dosage reduces expression at genes that negatively regulate components of the TLR4 signalling pathway, such as the MAPK cascade (Suppl. Fig. S1, Cluster III)^37,69,70^.

The acute inflammation group, where the signal was either substantially lower or higher in High-dose LPS compared to Low-dose LPS and PBS, consists of clusters IV and V. These enhancers are recognized as highly characteristic of the High-dose condition and associated acute inflammation. In Cluster IV, we once again see autophagy-related pathways. Gene promoters with substantially lower H3K27ac signal in High-dose cells were enriched in pathways for Il-6 and Il-8 production, which can have anti-inflammatory effects (Suppl. Fig. S1, Cluster IV)^69,70^. In Cluster V, we see many nucleotide-associated metabolic and catabolic processes, which have been shown to be increased with LPS stimulation in literature^71,72^. It is unclear why these processes are not increased in Low-dose cells.

The low-grade inflammation group, in which signal was either substantially increased or decreased in Low-dose LPS when compared to High-dose LPS or PBS, consists of clusters VI, VII, VIII, IX, and X. These enhancers are recognized as highly characteristic of the Low-dose condition and associated low-grade inflammation. Only Cluster VIII had any significantly enriched terms, most of which were related to histone and protein acetylation, primarily acetylation at lysines. LPS stimulation has been shown in literature to affect histone acetylation^73^, but we do not have enough information to determine which modifications (other than H3K27ac) are affected uniquely by Low-dose LPS, though aberrant histone acetylation has been found in multiple chronic inflammatory diseases^74,75^. However, the enhancers within this grouping were linked to some interesting genes within the immune system. For example, enhancers who had much higher signal in Low-dose were linked to genes like Csf1, Irf5, Rab11a, Ccr5, Irf7, Ticam2, Ifit3, and Irf1, while enhancers with lower signal in Low-dose were linked to genes like Arg1, Hdac5, and Treml1. Rab11a is responsible for transporting TLR4 from the endocytic recycling compartment to forming phagosomes^37^. This transport triggers the TRIF-dependent signalling pathway, which requires TRAM (Ticam2). TRIF-dependent signalling leads to increases in interferon-alpha and interferon-beta production which, in turn, activate interferon-induced genes such as Ifit3. Hdac5^76^ and Treml1^77^ both regulate the inflammatory response. Furthermore, reduction of Hdac5 expression was also associated with an increase of Irf1 and transcription of interferon-beta^76^. In addition, Arg1, which is a typical marker of anti-inflammatory macrophages^49^, is reduced in Low-dose while Irf5, shown to promote pro-inflammatory macrophage polarization^78^, is increased in Low-dose. Together, these suggest that, while the Low-dose cells are markedly different from High-dose cells, they do have a pro-inflammatory phenotype, which is consistent with previous studies^33^. It is interesting to note that Csf1, which is associated with anti-inflammatory macrophages, is highly increased in enhancer signal in Low-dose cells, however, previous studies have shown an association between Csf1 and chronic inflammation^79^.

Since enhancers are hotbeds of transcription factor binding activity, the enhancer regions were then scanned for transcription factor binding motifs that were separated into motifs uniquely and significantly enriched or diminished in low-grade inflammation, acute inflammation, or if the enrichment was correlated to LPS-dosage (Fig. 2c). Among the dosage-dependent affected motifs, we see multiple immune-related transcription factors that are increased with increasing LPS-stimulated inflammation such as JunB^80^, AP-1^81,82^, Atf4^83^, and NF-E2^51^.

In acute inflammation, we see several transcription factors that are important in the TLR-signaling pathway such as Nfil3^84,85^, Hoxa2^86^, Nrf2^87^, CEBP:AP1^88,89^, and OCT2^90,91^, all of which are activated with LPS stimulation in literature. In fact, the differential enrichment of Nrf2 between High-dose and Low-dose is also supported by previous research. Nrf2 is activated by LPS stimulation via the reduction of the protein Keap1, however, Keap1 protein has been shown to accumulate in Low-dose conditions^33,87^. The Mef2 family (Mef2a, Mef2b, Mef2c, Mef2d) of transcription factors motifs are significantly deficient in High-dose cells, but are present in Low-dose cells. It has been show that the Mef2 family is initially upregulated by LPS-stimulation but are soon downregulated^51^. It is possible that, in the Low-dose condition, the LPS-stimulation is not enough to lead to the downregulation of the Mef2 family.

Low-grade inflammation leads to fewer enriched motifs than deficient motifs. Motifs that are enriched only in Low-dose are ISRE, Irf2, and Klf4. Irf2^92,93^ and Klf4^94^ are both inflammation regulators while ISRE is an IFN-I stimulating response element which, when bound, activates genes in the inflammation pathway^95^. This is consistent with previously discussed upregulated enhancer-linked genes such as Irf7. Additionally, Irf2 has been shown to positively regulate the non-canonical inflammasome pathway, which, in turn, leads to increased GSDMD expression^96^. In fact, we do see increased GSDMD expression in our enhancer data, where it is located in Cluster VI. Low-dose also has reduced p65 motifs, a NF-κB subunit that is part of the canonical pathway involved in inflammation^97^. However, in monocytic cells that have already been stimulated with a gram-negative bacteria, a second stimulation shows reduced p65 activity^98,99^. There is a possibility that sustained low-dosage of LPS may lead to such a reaction. Furthermore, it appears that reduced p65 expression can be somewhat compensated for^100^.

We analyzed RNA-sequencing data to further understand the effect of LPS on murine immune cells. We see a similar pattern of RNA-seq data correlation as was in the ChIP-seq data (Fig. 3a). Average replicate correlation was 0.996 and the correlation between PBS and low-dose was similarly high (0.987). The high-dose replicates had approximately similar correlations with either PBS (0.892) or low-dose (0.914). Unlike the H3K27ac signal, there was not a decrease in RNA signal with increasing LPS-dosage (Suppl. Fig. S2a). Much like the differential peaks, the number of differentially expressed genes (DEGs) was lowest between PBS and Low-dose (155) with much greater differences between PBS and High-dose (3,249) closely followed by Low-dose and High-dose (3,061) (Suppl. Fig. S2b, Suppl. Tables S5 and S6). However, differentially expressed genes were largely equally split into up- and down-regulated genes for each comparison, except between PBS and Low-dose, where there were more genes upregulated in Low-dose than PBS samples. Many genes (~63%) that are differentially expressed in one comparison, are differentially expressed in multiple comparisons (Suppl. Fig. S2c).

**Figure 3.**
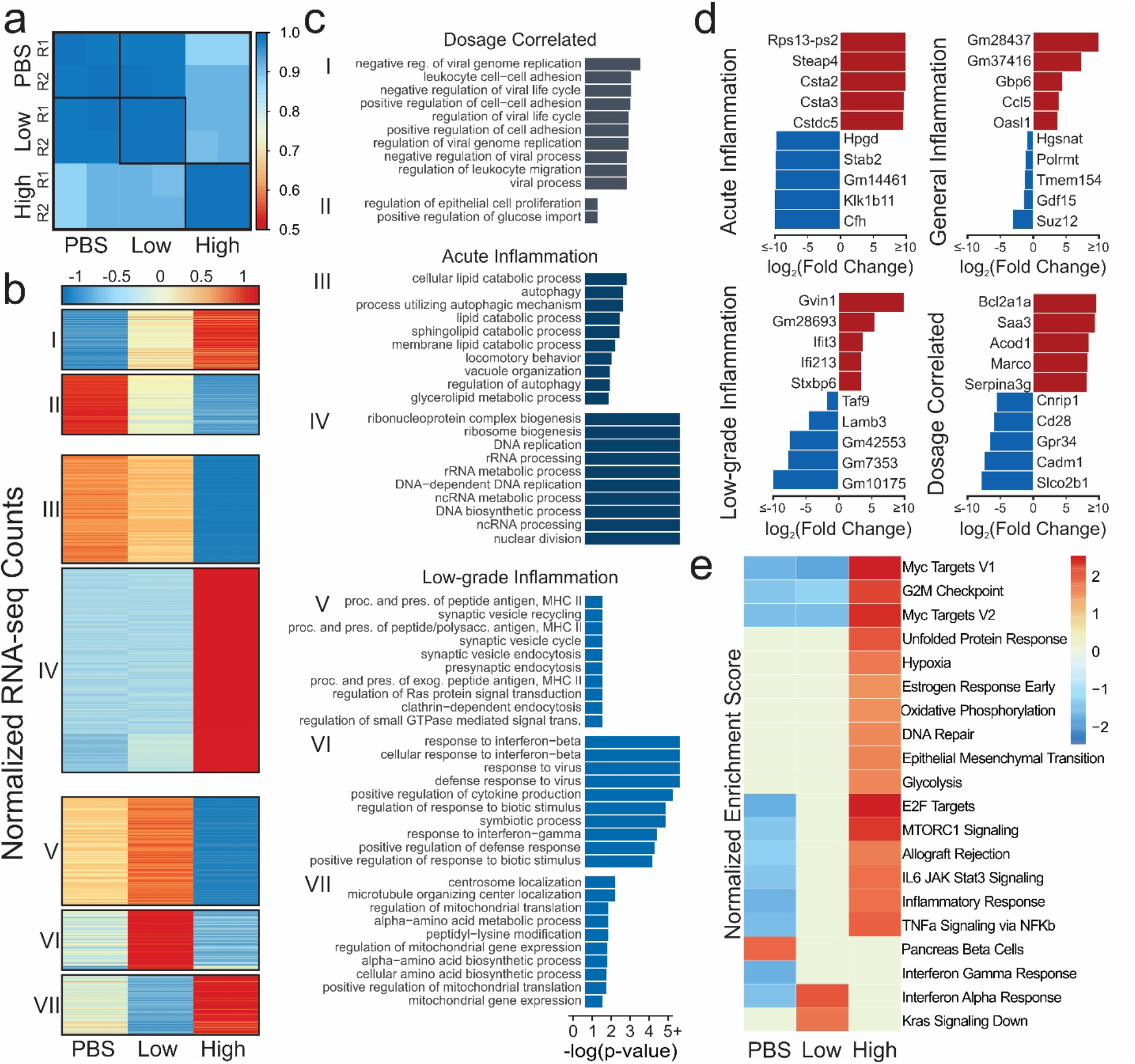
Effect of LPS dosage on gene expression. (a) Pearson’s correlation of normalized RNA-seq counts at genes (b) Heatmap of normalized gene expression of DEGs present in more than one comparison. (c) Significant biological process gene ontologies for genes that are differentially expressed in more than one comparison. (d) Top upregulated and downregulated genes for each condition. (e) Gene-set enrichment analysis. Color denotes normalized expression score if that pathway was significant (FDR < 0.05) in sample vs. rest.

The normalized RNA-seq expression data of DEGs were visualized in a heatmap (Fig. 3b). Each of these clusters were analyzed for enrichment of Gene Ontology biological process gene sets (Fig. 3c, Suppl. Table S7). Dosage correlated Clusters I and II, which increased or decreased with added LPS-dosage, are consistent with enhancer analysis. Both analyses identified leukocyte-related pathways were increased with increasing dosage. In addition, negative viral process was also increased with additional LPS. While LPS is from a bacteria, TLR4 can also be activated by viral ligands, thus there is large overlap in the genes in each of these ontologies^101^. We also see an reduction of glucose import with increased LPS-dosage, which is also consistent with literature^102^. Clusters specific to acute inflammation, Clusters III and IV, had similar autophagy and nucleic acid-related processes as the corresponding enhancer clusters. However, Cluster III also had more lipid metabolic processes that were down-regulated in High-dose conditions, which is consistent with previous research showing LPS-stimulation to decrease lipid catabolism^103^.

Finally, Clusters V, VI, and VII were specific to Low-dose. In Clusters V and VI, we see ontologies that are significantly upregulated in Low-dose compared to High-dose or PBS. These include MHC II antigen processing and response to interferon-beta. MHC II antigen processing is the means by which an exogenous antigen is prepared and presented on the cell surface for the activation of CD4+ T cells. These cell surface molecules that present the antigen are only upregulated in the TRIF-dependent pathway. Furthermore, what the macrophage secretes can also affect CD4+ T cell function and aberrant CD4+ T cells have been implicated in inflammatory diseases^104,105^. Interferon-beta related genes and motifs were also found to be enriched or have increased enhancer expression in Low-dose cells compared to High-dose or PBS. This suggests that epigenetics play a notable role in the etiology of low-grade inflammation. This is further supported by the peptidyl-lysine modification process in Cluster VII, which is down in Low-dose and high in High-dose. With closer inspection, Hdac2 and Hdac4 are included within this clusters. Hdac2 is considered to be crucial to the LPS inflammatory response while also mediating it, and decreased Hdac2 levels have been found in COPD patients^106–108^. Hdac4 is necessary for LPS-stimulated production of pro-inflammatory cytokines, and degradation leads to secretion of HMGB1, which is believed to play a significant role in sepsis^109,110^.

The fold-changes of the top genes significantly expressed or repressed in Acute Inflammation (High-dose vs. rest), Low-grade inflammation (Low-dose vs. rest), General Inflammation (PBS vs. rest), and Dosage-specific (increasing or decreasing from PBS to High-dose) were inspected (Fig. 3d). In each, we see some genes that have been pointed out in literature before as affected by inflammation or LPS in some way, such as Steap4^111^, Bcl2a1a^112^, Saa3^113^, Marco^114^, CFH^115^, Ifit3^116^, Lrrc14b^117^, Ccl5^118^, and CD28^119^. We also see multiple predicted genes that might be worth further *in vivo* or *in vitro* functional investigation.

Gene-set enrichment analysis was then performed using each of the conditions compared to the rest (Fig. 3e). The first aspect of note is that High-dose is significantly enriching many pathways that have been found to be enriched by LPS-stimulation in literature such as Myc^120,121^, hypoxia^122^, glycolysis^123^, and unfolded protein response^124^. We also see oxidative phosphorylation, which is generally believed to be reduced in LPS-stimulation as the cell transfers to glycolysis, though there is some conflicting evidence^125^. Yet, there are two reasons that it might be enriched in High-dose. First, oxidative phosphorylation is necessary for inflammatory resolution^126^. Or, it could be as a result of glucose starvation, which can cause cells to shift back towards oxidative phosphorylation^127^. In addition, there are also pathways that seem to be somewhat dosage-dependent, such as E2F targets^128^, MTORC1^129^, IL6 JAK Stat3 signaling^130,131^, and TNF-α^132,133^ that are hallmarks of inflammation. However, it is interesting to note that it appears low-dose cells are not undergoing as much replication, as seen in the reduced G2M checkpoint and glycolysis. Furthermore, it appears that, while slight interferon-gamma enrichment^134,135^ occurs in both Low-dose and High-dose (seen by deficiency in PBS), that low-dose has stronger enrichment of interferon-alpha and Kras Signaling down. Interestingly, interferon-alpha overexpression, which studies suggest can lead to chronic inflammation^136^, is a characteristic of systemic lupus eruthematosus, an autoimmune disease^137,138^. The reduction in Kras Signaling, which is part of the Ras/MAPK pathway, would also point to reductions in the cell-cycle of Low-dose cells^139^.

As differential transcript usage (DTU), such as alternative splicing, has been shown to be affected in the macrophage inflammatory response^140^, we also chose to examine DTU among the three experimental conditions as differential transcript usage can effect the function of the genes involved. There were a total of 309 genes that had significant DTU with predicted functional consequences across the three comparisons (PBS vs. Low-dose = 82, PBS vs. High-dose = 171, Low-dose vs. High-dose = 172) (Suppl. Fig. S3a). Genes with DTUs in PBS versus High-dose were entirely associated with RNA-splicing ontologies (data not shown), as well as many in Low-dose vs High-dose (Suppl. Fig. S3b). However, Low-dose versus High-dose did have several immune related ontologies, suggesting that DTU may play a role in the differing responses. In particular, we see response to interferon-beta and one of the DTU genes that was involved, IFI204, plays a key role in interferon-beta release^141^. We also found that, of the 172 DTUs in Low-dose vs. High-dose, 111 had (FDR<0.05) differential H3K27ac signal at their promoters (expected = 40) and 38 were linked to enhancers (expected = 22) for a total of 121 DTUs associated epigenetic regulation. This further suggests that epigenetic regulators may be playing a role in the differences between acute and low-grade inflammation.

The increase that we see of type I interferons-alpha and beta-related processes in Low-dose cells, both epigenetically and transcriptomically, as well as increased epigenetic enhancement of TRAM (RNA-seq FDR < 0.05, FC ≈ 1.8) led us to postulate that low-dose LPS might lead to more use of the TRIF-dependent pathway (which requires TRAM), while high-dose might lead to more use of the MyD88-dependent pathway (Fig. 2a and Fig. 3c,e)^37^. In order to better understand the mechanistic pathways that might be differentially affected by low-dosage of LPS, we performed additional experiments using TRAM-deficient and IRAK-M-deficient BMDMs (Fig. 4a). TRAM-deficient cells are incapable of utilizing the TRIF-dependent pathway and IRAK-M-deficient cells have an unregulated MyD88-dependent pathway. We see that WT-high-dose cells correlate the strongest with TRAM-deficient-high (r = 0.961) followed by the low-dose and PBS conditions (r = 0.866 – 0.915), while the two IRAK-M samples had the lowest correlation (High = 0.815, PBS = 0.762). Average correlation between the WT and TRAM-deficient low-dose and PBS samples was very high at r = 0.986. IRAK-M-deficient PBS was most highly correlated with the other two PBS samples, but IRAK-M-deficient high-dose had lower correlation with all samples (r = 0.809 – 0.904). When we perform principle component analysis of the most variable genes, we see fairly similar results (Fig. 4b). Due to the significant differences in IRAK-M-deficient cells, we chose to limit their inclusion in further analysis.

**Figure 4.**
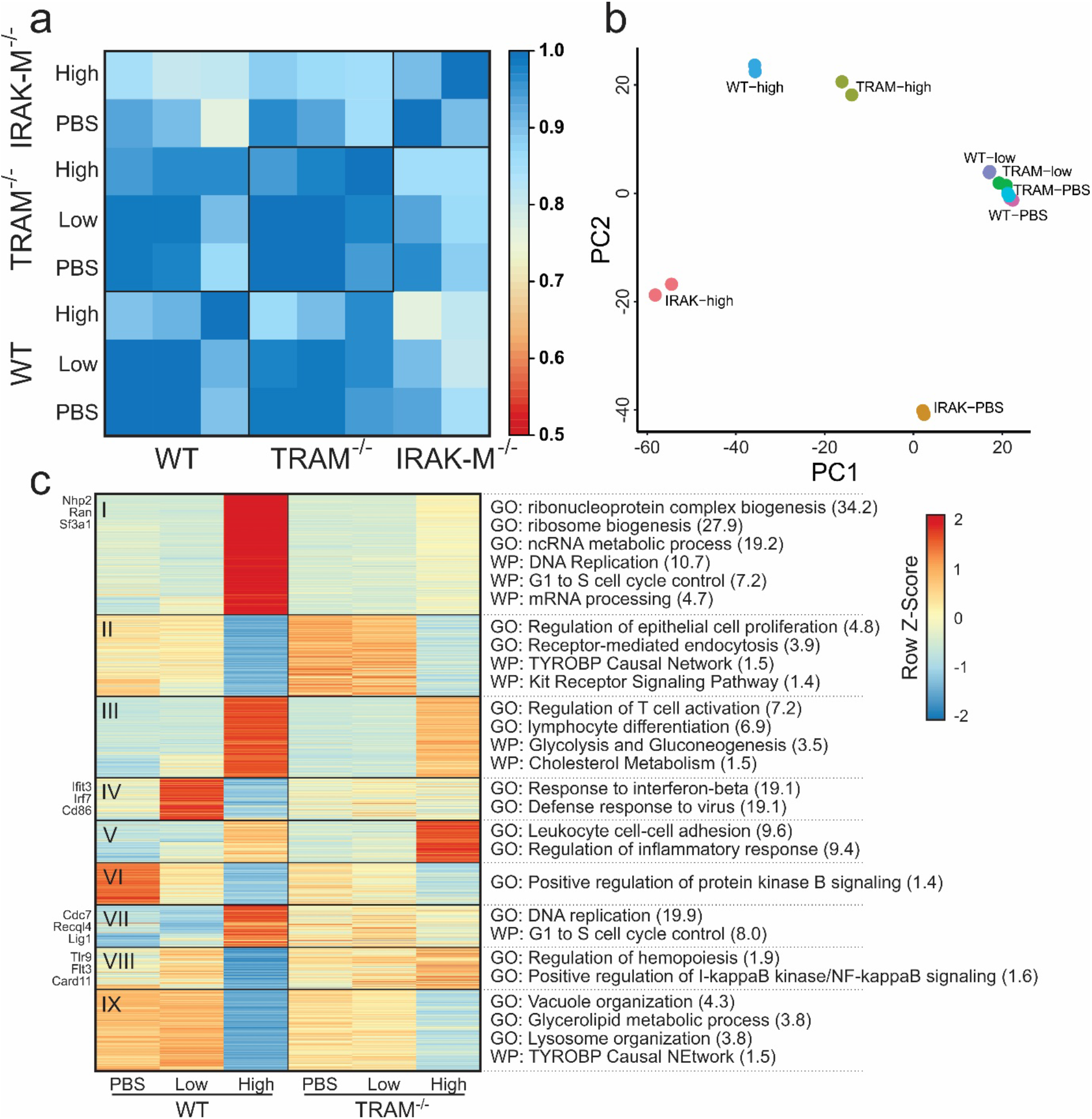
Effect of TLR4 pathway on LPS-dosage response (a) Pearson’s correlation of RNA-seq signal between conditions. (b) Principle Component Analysis of replicates using top 500 variable genes. (c) Heatmap of WT and TRAM-deficient gene express at varied LPS-dosage using DEGs from the WT comparisons. Significant pathways on right are from Gene Ontology or WikiPathways. Log(FDR) for each in parentheses.

We then analyzed gene expression across the three dosing conditions for both WT and TRAM-deficient cells, though we restricted the analysis to genes that were differential expressed between the WT conditions (Fig. 4c, Suppl. Tables S8 and S9). A decent number of genes were moderately affected by TRAM-deletion, but most maintained a similar expression pattern to WT. Despite this, there are 4 clusters of interest (Clusters I, IV, VII, and VIII) that stand out. Clusters I and VII both show a failure to upregulate DNA replication pathways in TRAM-deficient high-dose cells. While there is evidence of macrophage renewal due to inflammatory insult, it is not well understood, but it is possible that it is TRIF-dependent^142,143^.

On the other hand, Clusters IV and VIII both have a pattern of low-dose having the highest expression and high-dose the lowest in WT. There appears to be little to no change in TRAM-deficient cells. In Cluster VIII, TLR9 is an intracellular toll-like receptor that recognizes unmethylated CpG motifs in bacterial or viral DNA^144^. However, TLR9 has been shown to occur after LPS stimulation^145^ and TLR9 inhibition helps suppress excessive inflammation in bacterial sepsis^146–148^. Furthermore, it helps regulate antigen presentation in macrophages^149^ and participates in interferon-alpha production through IκB kinase signaling^150^. Interestingly, TLR9 is found to be activated through the NF-κB, Erk, Jnk, and p38 Mapk pathways which are not TRIF-dependent, so it is unclear why there is not an increase in TRAM-deficient cells as well^145,151^.

Another critical gene is Irf7, which is upregulated in WT-low but not in TRAM-low, that is important for the production of type I interferons^152^. Furthermore, Irf7 expression can lead to a feed-forward loop of type I interferon production, much like we see in WT-low^153,154^. However, research has shown that Irf7 is necessary for the Il-1β and the type I interferon response elicited through TLR4, and that Irf7 is induced through the TRIF-dependent pathway, consistent with the lack of Irf7 increase in TRAM-deficient cells. Yet, Irf7 is not overwhelming in WT-high cells, so there must an additional factor in low-dose conditions. Trim28 is a negative regulator of Irf7^152^ and is significantly reduced in WT-low compared to WT-high cells (FDR < 0.05, FC < 2), while remaining unchanged across the TRAM-deficient cells. If Trim28 is phosphorylated at serine 473 through a PKR/p38 MAPK/MSK1 signaling cascade, it is no longer capable of inhibiting Irf7^155^. Although knockdown of activated MSK1 does increase production of some inflammatory cytokines in neuroinflammation, Trim28 is responsible for regulating a staggering number of genes, so it is unclear if it is involved^155,156^. In addition, phosphorylation of Trim28 is not necessarily equivalent to reduced gene expression. Foxo3^157^ and Cflar^158^ have also been shown to inhibit Irf7 but neither had notable differences in gene expression.

## Discussion

Our genome-wide study of gene expression and histone modification changes due to LPS-dosage suggest that a low dosage of LPS leads to cells preferentially utilizing the TRAM/TRIF-dependent TLR4 signaling pathway. First, we found that these changes are seen in the variation of enhancers, the regulators of gene expression. Enhancers with increased signal in Low-dose samples were linked to genes related to the TRAM/TRIF-dependent pathway. In addition, promoters and motifs enriched in Low-dose samples were also associated with the interferon-beta response. Second, we showed that genes with increased expression only in Low-dose LPS, such as Stat1, Irgm1, Ifit3, and BST2, were significantly involved in interferon-beta and interferon-alpha responses. Furthermore, genes with differential transcript usage between Low-dose and High-dose were also associated with interferon-beta response. Finally, we compared the gene expression of WT samples to TRAM-deficient mice to determine that TRAM-deficiency abrogates the interferon-beta associated genes, indicating that the Low-dose increases in interferon-beta specifically rely on TLR4-associated TRAM/TRIF-dependent pathway, as opposed to MyD88 pathway. This preference for the TRIF-dependent pathway also prevented Low-dose samples to not alter nucleotide replication and metabolic pathways that were changed in High-dose samples preferentially going through the MyD88 pathway.

There is increasing appreciation for signal-strength dependent programming of both innate and adaptive immune systems, enabling complex and dynamic host responses to changing landscapes of infectious and inflammatory conditions^159,160^. Although much progress has been made with the dynamic programming of T helper cells exhibiting multi-staged activation and exhaustion under signal-strength and history dependent challenges^161^, the similar scenario of innate immune cell adaptation is still less understood. Due to limited systems approaches, even the most well-known concept of endotoxin tolerance regarding innate cell adaptation to repeated LPS challenges fails to clarify the complex innate immune adaptation dynamics. Past studies regarding endotoxin tolerance overly focused on dampened gene expression of limited inflammatory mediators^32^, and failed to address augmented induction of diverse immune-; metabolic-; and proliferative related genes involved in complex adaptation to higher dosages of endotoxin. Much less is known about innate responses to pathologically relevant subclinical low dose LPS, highly prevalent in humans with chronic conditions due to mild leakage of mucosal barriers^162^. The lack of systems and clear understanding of signal strength and history-dependent adaptation to LPS underlies our limited translational success in treating related diseases ranging from acute sepsis to chronic cardiovascular diseases. Our current study provides a comprehensive assessment of gene expression dynamics as well as corresponding epigenetic variations in monocytes challenged with rising dosages of LPS.

Confirming limited previous studies, our collected data reveal that higher doses of LPS not only cause suppression of certain subsets of inflammatory genes, but also potently induce wide arrays of genes involved in altered immune metabolism and proliferative potential. Further characterization of these altered gene expression landscape may help better explain the compound phenotypes of pathogenic inflammation and immune-suppression observed in septic leukocytes collected from human sepsis patients and model murine septic animals^163,164^. In contrast, low dose LPS preferentially induces inflammatory interferon responsive genes, recently shown to be expressed in inflammatory monocytes collected in vivo from various chronic inflammatory disease models including lupus and atherosclerosis^138,165,166^.

Our integrated analysis of epigenomic changes at enhancer and promoter regions complements our gene expression data, in further revealing the preferential usage of TRAM/TRIF pathway by low dose LPS. Our finding is consistent with limited previous studies showing that the TRAM/TRIF-dependent pathway is favored in Low-dose LPS conditions and critical for lesion development in atherosclerosis^33^. In addition, we identified signature transcription factors involved in monocyte activation by Low-dose LPS such as IRF1, 5 and 7, with IRF5 previously reported to be involved in monocyte priming by Low-dose LPS^167^. The preferential enrichment of additional transcription factors such as IRF2 and KLF4 is also interesting, and may provide additional insight regarding the regulation of low-grade inflammatory monocytes. Low-dose LPS also enhanced the expression of Rab11a, a molecule involved in endocytic recycling of TLR4, providing a further mechanistic explanation for our previous observation that internalization of TLR4-LPS complex and the activation of TRAM/TRIF pathway are required for sensing Low-dose LPS^35^.

Collectively, our integrative systems study further clarifies the highly complex and dynamic adaptation of macrophages to rising dosages of LPS, and reveals more of the inner workings of underlying mechanisms, yet much more research will be required to fully understand how immune pathways and components interact in the dynamic ontogeny of macrophage activation states related to the etiology of low-grade inflammation. Studies that utilize time courses, additional LPS concentrations, and other transgenic mice all would be beneficial for truly unveiling low-grade inflammation. Additional epigenomic studies may also reveal the causal relationships at play and possible therapeutic targets. Together, this could lead to identification of relevant molecular targets in human immune cells for future clinical applications.

## Methods

### Mice

C57/BL6 mice purchased from the Jackson Laboratory were bred and maintained in pathogenic-free conditions. Male 8-12 week old mice were used in this study. The Institutional Animal Care and Use Committee (IACUC) approved all procedures performed on the mice.

### Cell Culture

Crude BM cells were isolated from the mice and cultured as previously published^33^. Briefly, cells were cultured in RPMI complete media (RPMI 1640 with 10% FBS, 2 mM L-glutamine, and 1% penicillin/streptomycin) along with M-CSF (10 ng/mL) and either PBS, low-dose LPS (100 pg/mL), or high-dose LPS (1 μg/mL). Fresh LPS was added every two days and the cells were harvested after 5 days.

### Chromatin Shearing

Samples containing 10^6^ cells were centrifuged at 1600 g for 5 minutes at 4°C. Each sample was washed twice with cold 1 mL PBS and resuspended in 9.375mL of PBS. Then, 0.625 mL of 16% formaldehyde was added and the samples were incubated on a shaker for 5 minutes at room temperature. The samples were then quenched with 0.667 mL of 2M glycine and incubated for 5 minutes at room temperature on a rotator. The cells were then centrifuged at 1600 g for 5 minutes and washed twice with 1 mL cold PBS. The pellet was resuspended in 130 μL of Covaris sonication buffer (10 mM Tris-HCl, pH 8.0, 1 mM EDTA, 0.1% SDS and 1x protease inhibitor cocktail (PIC)) and sonicated with a Covaris S220 sonicator using 75 W peak incident power, 5% duty factor, and 200 cycles per burst for 16 minutes at 4°C. Sonicated samples were centrifuged at 16,100 g for 10 minutes at 4°C before the supernatant containing sheared chromatin was removed to a fresh tube. 2.4 μL of sheared chromatin was then mixed with 46.6 μL of IP buffer (20mM Tris-HCl, pH 8.0, 140 mM NaCl, 1 mM EDTA, 0.5 mM EGTA, 0.1% (w/v) sodium deoxycholate, 0.1% SDS, 1% (v/v) Triton X-100, 1% freshly added PMSF and PIC) to generate a 50μL sample containing chromatin from 20,000 cells for MOWChIP-seq.

### MOWChIP-seq

Sonicated chromatin samples of 20,000 cells per assay were profiled for H3K27ac (abcam, cat: ab4729, lot: GR312651-2) with MOWChIP-seq as described in our previous publications^53,58^. Libraries for sequencing were prepared using the Accel-NGS 2S Plus DNA Library Kit (Swift-Bio) and samples were sequenced using an Illumina HiSeq 4000 with single-end 50 nt reads.

### RNA-seq

10,000 cells were used to produce each RNA-seq library, with two replicates for each genotype and experimental condition. RNA was extracted into a 30-μL volume using the RNeasy Mini Kit (74104, Qiagen) and RNase-Free DNase Set (79254, Qiagen), following the manufacturer’s instruction. The extracted mRNA was then concentrated by ethanol precipitation and resuspended in 4.6 μL of RNase-free water. Next, we used the SMART-seq2 protocol^55^, with minor modifications, to prepare cDNA. 2 μL of oligo-dT primer (100 μM) and 2 μL of dNTP mix (10 mM) were added to 2 ng of mRNA in 4.6 μL of water. The mRNA solution was denatured at 72 °C for 3 min, then immediately placed on ice. Next, 11.4 μL of reverse transcription mix [1 μL of SuperScript II reverse transcriptase (200 U/μL), 0.5 μL of RNAse inhibitor (40 U/μL), 4 μL of Superscript II first-strand buffer, 1 μL of DTT (100mM), 4 μL of 5 M Betaine, 0.12 μL of 1 M MgCl_2_, 0.2 μL of TSO (100 μM), 0.58 μL of nuclease-free water] was added to the mRNA solution. For the reverse transcription reaction, the solution was incubated at 42 °C for 90 min, followed by 10 cycles of 50 °C for 2 min, 42 °C for 2 min, then inactivation at 70 °C for 15 min. 20 μL of the resulting solution (first-strand mixture) was then mixed with 30 μL of PCR mix [25 μL KAPA HiFi HotStart ReadyMix, 0.5 μL IS PCR primers (10 μM), 0.5 μL Evagreen dye, and 4 μL nuclease-free water] and amplified using the program 98 °C for 1 min, followed by 9-11 cycles of 98 °C 15 s, 67 °C 30 s, 72 °C 6 min. Finally, the cDNA was purified using 50 μL of SPRIselect beads. RNA-seq libraries were generated with the Nextera XT DNA Library Preparation kit (FC-131-1024, Illumina) and manufacturer’s protocol, using approximately 600 pg of purified cDNA from each sample. Samples were sequenced using an Illumina HiSeq 4000 with single-end 50 nt reads.

### Data Processing

Unless otherwise mentioned, all data analysis was performed with Bash scripts or with R (The R Foundation) scripts in RStudio. Sequencing reads were trimmed using default settings by Trim Galore! (Babraham Institute). Trimmed reads were aligned to the mm10 genome with Bowtie^168^. Peaks were called using MACS2 (q < 0.05)^169^. Blacklisted regions in mm10 as defined by ENCODE were removed to improve data quality^170^. Mapped reads from ChIP and input samples were extended by 100 bp on either side (250bp total) and a normalized signal was calculated.

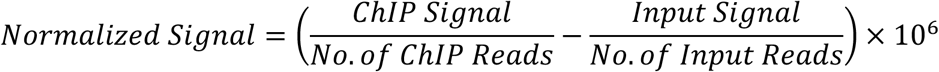

For Pearson’s correlation, the signal was calculated around the promoter region (TSS +/− 2kb) and plotted with the corr and levelplot functions. For visualization in IGV (Broad Institute), the signal was calculated in 100bp windows over the entire genome and output as a bigWig file. RNA-seq data was quantified using Salmon^171^ against the mm10 transcriptome using a full decoy and normalized counts were calculated with DESeq2^172^.

### Enhancers Analysis

To call enhancers, we considered H3K27ac^high^ regions that did not intersect with promoter regions to be enhancer regions. First, consensus H3K27ac peak sets were generated for each of the experimental conditions. Peak widths were expanded to be 1000 bp long (summit +/−500bp). Promoters were defined as TSS +/− 2000 bp. Any H3K27ac 1kb regions that intersected with a promoter region was removed and the remaining regions were designated as putative enhancers. The signal at each of the putative enhancers was then correlated (Spearman) to RNA-seq gene expression values within the same topological domain. Putative enhancers were linked to the gene with the highest correlation, however, links were only considered significant if the Spearman correlation coefficient (SCC) > 0.25 and if the correlation was considered significant if both empirical and quantitative p-values were less than 0.05. For both p-values, the SCC was calculated between the given putative enhancer and all genes on the same chromosome. The empirical p-value was then calculated as the fraction of genes on the same chromosome that has a higher correlation than the currently linked gene. The quantitative p-value was calculated by treating the calculated SCC values as a distribution and using the R function pnorm to calculate a significance. Motif analysis was performed to determine enriched transcription factor binding motifs among the enhancer regions with HOMER^173^ (with options –size 1000 –mask –p 16 –nomotif).

### RNA-seq Analysis

Differential gene expression analysis was performed using DESeq2, where genes with a fold-change >= 2 and FDR < 0.05 were considered to be significantly differentially expressed. Boxplots and MA plots were done in R using ggplot and ggpubr. Clustering was performed using clusterProfiler. Gene-set enrichment analysis was performed with GSEA^174,175^ using the Hallmark gene set and gene-set level permutation. Gene sets were considered significant if the FDR < 0.05. Significant differential transcript usage (p < 0.05, dIF > 0.1) was determined using IsoformSwitchAnalyzeR^176,177^ with the default DEXSeq^178^. Data output was then ran through CPAT^179^, PFAM^180^, SignalP^181^, NetSurfP-2.0^182^, and results were combined back into IsoformSwitchAnalyzeR to determine genes that might have functional consequences as a result of the DTUs. Genes were analyzed for gene ontologies with clusterProfiler.

## Data Availability

The ChIP-seq and RNA-seq data sets are deposited in the Gene Expression Omnibus (GEO) repository with the following accession number: GSE168190.

## Acknowledgments

This work was supported by US National Institutes of Health grants R01EB017235 (C.L.) and R01AI136386 (L.L.), and Virginia Tech ICTAS Center for Engineered Health seed grant (C.L.).

## Author contributions

C.L. and L.L. designed the experiments and supervised the research. S.G. conducted the animal experiments and produced the cell samples. Y.P.H. and Z.Z. conducted the ChIP-seq and RNA-seq assays. L.B.N. analyzed the data. L.B.N., Y.P.H., S.G., L.L. and C.L. wrote the manuscript. All authors discussed the results and commented on the manuscript prior to submission.

