## Supplementary Figures and Tables for "Epigenomic and transcriptomic differences between low-grade and acute inflammation in LPS-induced murine immune cells"

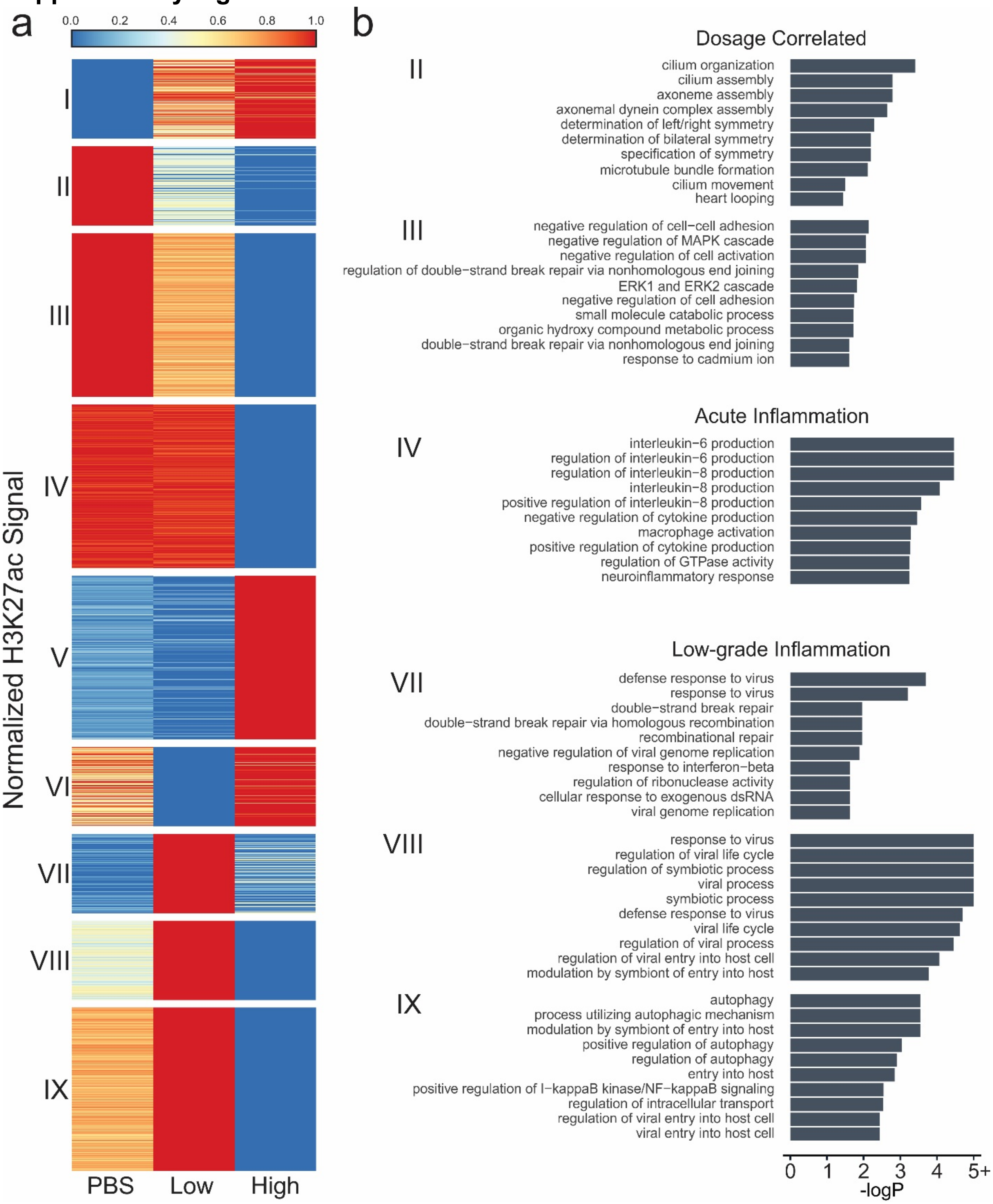


**Supplemental Figure S1** H3K27ac signal at promoters for murine BMDMs dosed with PBS, low-dose LPS, or high-dose LPS. (a) Normalized H3K27ac signal around promoter regions (TSS +/- 2 kb) with differential marking (FDR < 0.05, fold-change >= 2) clustered using k-means clustering. (b) Gene ontologies that are significantly enriched (FDR < 0.05) in the associated clusters from (a).

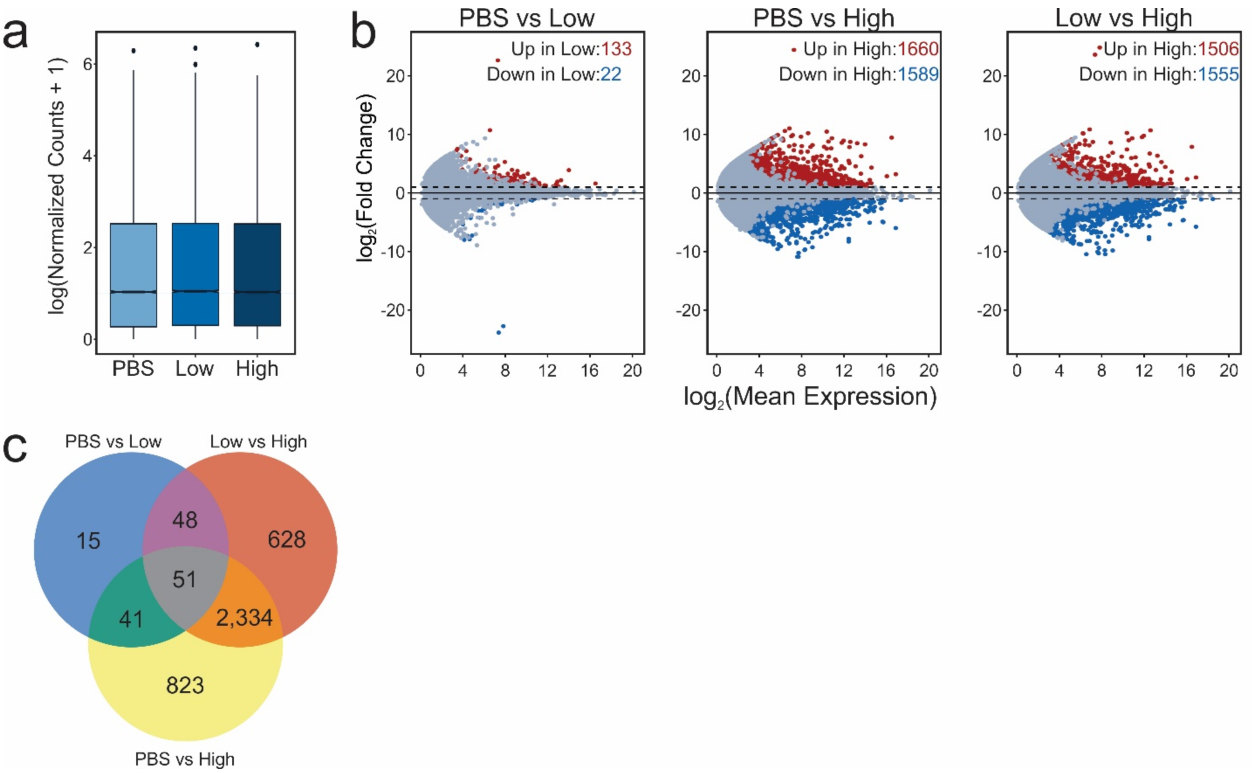


**Supplementary Figure S2** Overview of gene expression changes due to LPS. (a) Boxplots of normalized RNA-seq counts at genes (b) MA-plots of the fold change between two samples of a comparison versus the mean gene expression of that gene. Color denotes significance (FDR < 0.05, FC >= 2) (c) Overlap of differentially expressed genes.

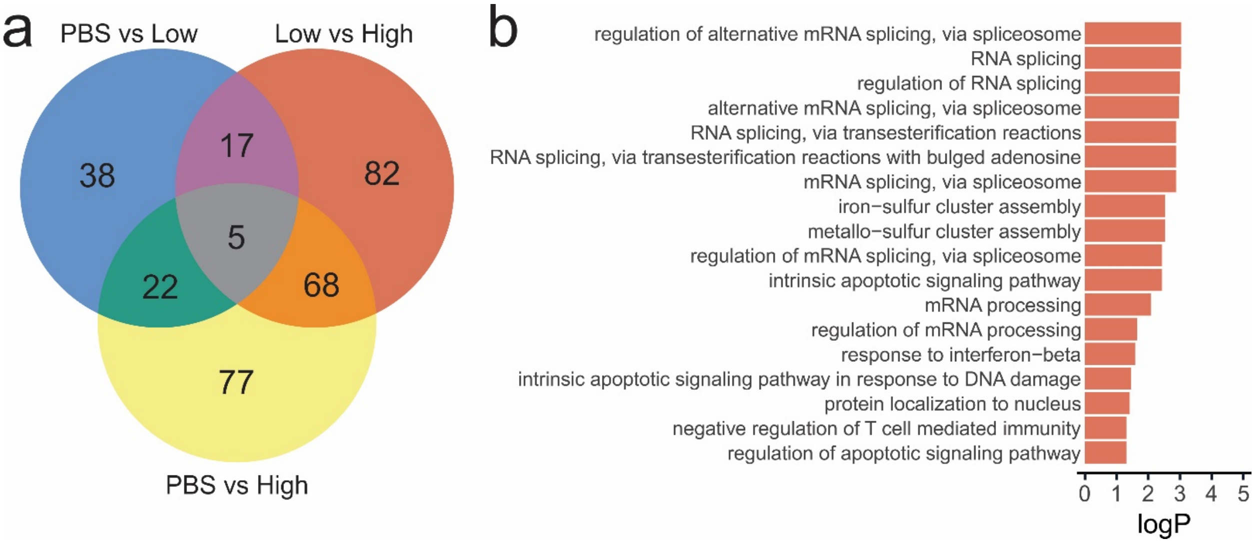


**Supplementary Figure S3** Effect of LPS on differential transcript usage (a) Overlap of genes with significant differential transcript usage among the three conditions. (b) Gene ontologies significantly overrepresented in genes with DTUs in High-dose compared to Low-dose.

**Supplementary Tables**

Table S1: WT H3K27ac Metadata

| Sample | Total Reads (millions) | Trimmed Reads (millions) | Aligned Reads (millions) | Alignment (%) | Peaks | Consensus Peaks |
|---|---|---|---|---|---|---|
| PBS – 1 | 13.8 | 13.8 | 12.0 | 87.1 | 34,372 | 25,679 |
| PBS – 2 | 15.9 | 15.9 | 14.0 | 88.0 | 46,067 | |
| Low – 1 | 21.4 | 21.3 | 18.2 | 85.4 | 34,967 | 21,597 |
| Low – 2 | 19.4 | 19.4 | 16.8 | 86.8 | 28,091 | |
| High – 1 | 19.4 | 19.4 | 16.6 | 83.5 | 20,685 | 14,659 |
| High – 2 | 21.0 | 21.0 | 18.1 | 86.2 | 23,775 | |

Table S2: RNA-seq Metadata

| Genotype | Sample | Total Reads (millions) | Trimmed Reads (millions) | Aligned Reads (millions) | Alignment (%) |
|---|---|---|---|---|---|
| WT | PBS-1 | 18.0 | 18.0 | 15.0 | 83.4 |
| | PBS-2 | 16.9 | 16.9 | 14.3 | 84.7 |
| | Low-1 | 15.6 | 15.6 | 12.9 | 83.2 |
| | Low-2 | 15.1 | 15.1 | 12.6 | 83.8 |
| | High-1 | 16.8 | 16.8 | 14.2 | 84.5 |
| | High-2 | 14.0 | 14.0 | 11.9 | 85.3 |
| $TRAM^{-/-}$ | PBS-1 | 13.7 | 13.7 | 10.9 | 79.5 |
| | PBS-2 | 14.2 | 14.2 | 11.7 | 82.2 |
| | Low-1 | 14.0 | 14.0 | 11.3 | 80.7 |
| | Low-2 | 15.0 | 14.9 | 11.8 | 79.1 |
| | High-1 | 30.4 | 30.4 | 25.2 | 83.1 |
| | High-2 | 14.2 | 14.2 | 11.0 | 77.9 |
| $IRAK\text{-}M^{-/-}$ | PBS-1 | 13.4 | 13.4 | 11.1 | 82.7 |
| | PBS-2 | 17.7 | 17.7 | 14.8 | 83.8 |
| | High-1 | 16.2 | 16.2 | 13.8 | 85.4 |
| | High-2 | 18.3 | 18.3 | 14.7 | 80.7 |

Table S3: Clusters of gene-linked enhancers (in "Supplementary Table_S3.xlsx")

Table S4: Gene ontology of gene-linked enhancer clusters (in "Supplementary Table_S4.xlsx")

Table S5: Differentially expressed genes between WT experimental conditions (in "Supplementary Table_S5.xlsx")

Table S6: Clustered RNA-seq counts of genes differentially expressed in WT (in "Supplementary Table_S6.xlsx")

Table S7: Gene ontology of differentially expressed gene clusters (in "Supplementary Table_S7.xlsx")

Table S8: Clustered WT and $TRAM^{-/-}$ RNA-seq counts of genes differentially expressed in WT (in "Supplementary Table_S8.xlsx")

Table S9: Gene ontology of WT and $TRAM^{-/-}$ RNA-seq clusters (in "Supplementary Table_S9.xlsx")
